# Intracellular nanoparticle delivery by oncogenic KRAS-mediated macropinocytosis

**DOI:** 10.1101/658039

**Authors:** Xinquan Liu, Debadyuti Ghosh

## Abstract

The *RAS* family of oncogenes (*KRAS, HRAS, NRAS*) are the most frequent mutations in cancers and regulate key signaling pathways that drive tumor progression. As a result, drug delivery targeting *RAS*-driven tumors has been a long-standing challenge in cancer therapy. Mutant *RAS* activates cancer cells to actively take up nutrients, including glucose, lipids, and albumin, via macropinocytosis to fulfill their energetic requirements to survive and proliferate. Here, we exploit this mechanism to deliver nanoparticles in cancer cells harboring activating *KRAS* mutations. We have synthesized stable albumin nanoparticles that demonstrate significantly greater uptake in cancer cells with activating mutations of *KRAS* than monomeric albumin (i.e. dissociated form of clinically-used *nab-paclitaxel*). From pharmacological inhibition and semi-quantitative fluorescent microscopy studies, these nanoparticles exhibit significantly increased uptake in mutant *KRAS* cancer cells than wild-type *KRAS* cells by macropinocytosis. Importantly, we demonstrate that their uptake is driven by *KRAS*. This nanoparticle-based strategy targeting *RAS*-driven macropinocytosis is a facile approach towards improved delivery into *KRAS*-driven cancers.

## Background

Mutations of the *RAS* oncogenes (*HRAS, NRAS, and KRAS*) are the most frequent mutations in human cancers and are present in 25% of all cancers. Among the three isoforms of *RAS* genes, *KRAS* is the most frequent mutated (85% in all *RAS* driven cancers). In particular, hyperactivated mutations of *RAS* oncogenes initiate and drive tumor progression in a significant subset of lung, colorectal, and pancreatic cancers ^1^. Patients with oncogenic *RAS* mutations have poor prognosis in colorectal ^2^ and pancreatic cancers ^3^. As a result, drug delivery targeting RAS-driven tumors has been a long-standing goal for cancer therapy ^1,4^. However, targeting cancers with *RAS* mutations has been a significant challenge due to the poor therapeutic index of existing *RAS* inhibitors ^1^. Consequently, approaches that enhance delivery and accumulation of RAS-targeting therapeutics would greatly advance and significantly improve patient outcomes.

*RAS* hyperactivation drives cancer cell survival and proliferation by altering metabolic requirements of the cells to upregulate intracellular uptake; as a result, mutant *RAS* drives the uptake of numerous solutes. *RAS* proteins are GTPases that act as “molecular switches”, effectively cycling between binding to guanosine triphosphatase (GTP) and guanosine diphosphatase (GDP) ^5^. During homeostasis, *RAS* protein toggle between binding to GTP in its active state and GDP in its non-stimulated, inactive state. At rest, *RAS* protein is bound to GDP in its inactive state. Upon stimulation by growth factor cues, GDP is released and *RAS* binds to GTP, which subsequently activates downstream RAF/MEK/ERK signaling axis, resulting in cell proliferation. *RAS* activation also spurs on PI3K and RalGDS effectors, which also stimulate cell proliferation, migration, and survival ^6^. Then, GTPase activation protein stimulates breakdown of GTP via hydrolysis, producing GDP to bind and inactivate RAS. During tumorigenesis, specific mutations of *RAS* cause constitutive RAS-GTP binding and subsequent constitutive activation of downstream effectors resulting in uncontrollable cell proliferation and survival ^6^. Here, oncogenic *RAS* reprograms downstream signaling and alters cellular metabolism to fulfill the nutrient requirements of these actively proliferating cancer cells. RAS-transformed cells activate RAF/MEK/ERK signaling to increase glycolysis ^7,8^, non-oxidative pentose phosphate pathway ^7^, and hexosamine biosynthesis pathways (reviewed in ^9^), which increase biomass synthesis needed for cell survival and proliferation. In addition, cells have evolved to use available sources of lipids, proteins, and nutrients for cell survival and proliferation. RAS-driven cancers can scavenge nutrients intracellularly and extracellularly for their survival. Oncogenic *RAS* proteins stimulate macropinocytosis in quiescent fibroblasts ^10^ and cancer cells ^11–13^ to “drink” in surrounding bulk fluid and scavenger extracellular lipids and proteins. *HRAS* overexpressing embryonic fibroblasts and RAS-transformed cells demonstrate increased membrane ruffling characteristic of macropinocytosis and higher intracellular content of lysophospholipids ^10,14^. Macropinocytosis is a fluid-phase endocytic process whereby cells form membrane ruffles upon extracellular or intracellular cues, resulting in the formation of large diameter vacuoles (0.2 to 5 μm), or macropinosomes, that are able to transport solutes intracellularly ^15,16^. RAS-transformed cells use the macropinocytosis program to fulfill their metabolic dependency to maintain their growth and survival. Lipids, glutamine, and in particular albumin have been actively scavenged by RAS-transformed fibroblasts, breast and pancreas cancers harboring activating *RAS* mutations ^11–13,17,18^. Collectively, these findings strongly indicate that albumin is actively macropinoctyosed by RAS-transformed cells *in vivo* for metabolic needs. Towards this end, is it feasible to exploit this vulnerability of nutrient transport to deliver drug carriers?

Attempts have been tried to delivery therapeutic payloads encapsulated in exosomes and lipoprotein nanostructures to cancer cells via macropinocytosis ^19–21^. We hypothesize that albumin-based particles can exploit the macropinocytosis pathway of RAS-driven cancer cells for intracellular delivery. Albumin has been used as a carrier to deliver different drugs for various diseases including inflammation and cancer (reviewed in ^22^). In particular *nab-paclitaxel*, or albumin-complexed paclitaxel, has been shown in combination therapy to improve overall survival compared to monotherapy and is standard of care for the treatment of advanced pancreatic cancer ^23^. While albumin has been used, it has not been developed to actively explicitly used in mechanistic, macropinocytosis-driven delivery into mutant KRAS-specific cancers. Here, cross-linked albumin nanoparticles demonstrate enhanced uptake in oncogenic *RAS* cells compared to control cells with wild-type *RAS* by non-ligand mediated macropinocytosis. The physicochemical properties of the nanoparticles are tunable and they are colloidally and physiologically stable. Interestingly, these particles exhibit greater uptake than equivalent amounts of monomeric albumin (i.e. present in *in vivo* dissociated *nab-paclitaxel*). Through microscopy-based quantification, these NPs co-localize in macropinosomes. Through pharmacological inhibition and genetic knockdown experiments, we demonstrate that these NPs can be endocytosed via RAS-driven macropinocytosis. These collective findings demonstrate that the macropinocytosis pathway of oncogenic *RAS* cancer cells can be exploited for nanoparticle delivery. By understanding this mechanism between the specific cancer pathway and its impact on delivery, it will be feasible to develop drug carriers for pathway-specific, targeted delivery. This work has the impact to greatly improve upon drug delivery and targeting to RAS-driven cancers.

## Materials and Methods

### Synthesis of serum albumin nanoparticles

Nanoparticles were synthesized by modified desolvation methods [1]. Briefly, bovine serum albumin (BSA, Fraction V, Fisher Scientific) was dissolved in 10 mM sodium chloride solution to make a 1.5 % (w/v) BSA solution. The pH of the solution was adjusted to 9.0 with sodium hydroxide. The desolvation agent was a mixture of methanol and ethanol at the ratio of 7:3 (v/v). Then, 4 mL of the desolvation agent was added into 1 mL BSA solution using a syringe pump (KD Scientific) at 1 mL/min under constant stirring. Subsequently, 8% glutaraldehyde solution (Sigma-Aldrich) was added to the system to induce particle cross-linking. Cross-linking process was allowed under stirring at room temperature for 12 hours. The synthesized nanoparticles were washed with water for three times, using centrifugal filter membrane units (molecular weight cutoff 100 kDa, Amicon). Fluorescent fluorescein isothiocyanate (FITC, ThermoFisher Scientific) and Cyanine 7 (Cy7, Lumiprobe) was conjugated to monomeric BSA according to manufacturer’s protocol, respectively. Fluorescently labelled nanoparticles (FITC-NP and Cy7-NP) were synthesized by the same procedures as described above using FITC-BSA or Cy7-BSA instead of BSA.

### Characterization of nanoparticles

The hydrodynamic diameter and zeta potential of the synthesized nanoparticles were characterized using Zetasizer Nano ZS (Malvern) with 173° backscatter angle. The morphology of the nanoparticles was observed by transmission electron microscope (TEM). Nanoparticles solution was spread on a carbon coated grid and negatively stained with 2% uranyl acetate. The grid was air-dried and then observed by TEM (FEI Tecnai).

### Cell lines and cell culture

Human breast cancer cell lines, MDA-MB-231 harboring oncogenic *KRAS* mutation and MDA-MB-468 with wildtype *KRAS*, were purchased from American Type Culture Collection (ATCC). DMEM/high glucose medium (Corning) supplemented with 10 % fetal bovine serum (Gibco) and 100 U/mL penicillin-streptomycin (Gibco) was used to maintain both cell lines. Cells were kept in a humidified atmosphere with 5 % carbon dioxide at 37 C°.

### Measurement of intracellular uptake of nanoparticles by flow cytometry

MDA-MB-231 and MDA-MB-468 cells were seeded in 24-well plates at a density of 4×10^5^ cells/well, respectively. After attachment, cells were starved in serum free medium overnight. To compare the difference in uptake of monomeric albumin and nanoparticles, cells were respectively incubated with FITC-BSA and FITC-NP for 2 hours at different concentrations of 0.5 mg/mL, 1.0 mg/mL and 1.5 mg/mL (equivalent amount of albumin). To evaluate the inhibitory effect of macropinocytic inhibitor 5-(N-ethyl-N-isopropyl) amiloride (EIPA, Sigma-Aldrich) on the uptake of nanoparticles, cells were pre-treated with 25 μM, 50 μM and 75 μM EIPA for 30 minutes, respectively. Then, cells were incubated with 500 μg/mL FITC-NP for another 30 minutes. After each treatment, cells were placed on ice and washed with ice-cold PBS for three times. Cells were collected in PBS buffer and stained with propidium iodide (PI, Sigma-Aldrich). Samples were then analyzed by a flow cytometer (Accuri, BD Biosciences). PI-positive cells were excluded as dead cells.

### Measurement of macropinocytic index

Macropinocytotic index was measured by an image-based method with slight modification [2,3]. Cells were plated in a 24-well plate with a circular cover glass in each well. Cells were incubated with serum free medium overnight after reaching 60-70% confluency. The cells were incubated with 1 mg/mL TMR-dextran in serum free medium for 30 min. After treatment, cells were washed with ice-cold PBS for 5 times and fixed with 3.7% formaldehyde solution for 30 min at room temperature. DAPI solution was added to stain the nucleus of the cells. The cover glass was then placed cell side down onto a glass slide with a drop of mounting medium. Cell images were randomly captured using an Olympus IX-83 inverted fluorescence microscope with a 100× phase objective. A z-stack of frames throughout the entire height of cell monolayers was aquored. The z-stack images were then collapsed to a single image using extended focus imaging projection (CellSens 1.16). To calculate the macropinocytic index, the total cell area was first selected from phase contrast images using the polygon selection tool of ImageJ. Then the region of interest was applied onto the corresponding TMR-dextran image with thresholding for macropinosomes. The total area of macropinosomes was computed. The macropinocytic index was calculated as the following: macropinocytotic index = total area of macropinosomes/cell number.

### Colocalization of nanoparticles with macropinosomes

To quantify the colocalization of nanoparticle with macropinosomes, cells were plated in the same way as described in the measurement of macropinocytic index. Then cells were incubated with 1 mg/mL TMR-dextran and 1 mg/mL Cy7 labelled nanoparticles simultaneously. A group of cells were treated with 25 μM EIPA to evaluate the effect of macropinocytic inhibition on colocalization. After incubation, cells were fixed and sealed onto glass slides. Cell images were captured using an Olympus IX-83 inverted fluorescence microscope with a 100× phase objective. A z-stack of frames throughout the entire height of cell monolayers was acquired. The z-stack images were then collapsed to a single image using extended focus imaging projection. Background was subtracted with a constant of 1500. For each channel, the contrast was adjusted to the same scale. The Pearson correlation coefficient (PCC) between pixels of TMR-dextran and pixels of Cy7 nanoparticles were analyzed by Cellsense 1.16 (Olympus).

### Knockdown of KRAS protein expression and nanoparticle uptake

MDA-MB-231 cells were seeded in 6-well plates at a density of 1 × 10^6^ cells/well. After 24 hours, cells were transfected with siRNAs against *KRAS* gene (SMARTpool: Accell *KRAS* siRNA, Dharmacon) at a final concentration of 0.5 μM, 1.0 μM and 1.5 μM in Accell siRNA delivery media, respectively. A non-targeting siRNA was used as a negative control. 120 hours after transfection, cells were harvested for western blot analysis. Cells were lysed using RIPA buffer (Thermo Scientific) with protease inhibitor (Roche). The protein concentrations were determined by BCA protein assay reagent kit (Thermo Scientific). Equivalent amounts of lysates (20 μg total protein per lane) were loaded and separated by 10% SDS-PAGE (Invitrogen Bolt Bis-Tris Plus gel). Then, proteins were transferred onto a low fluorescence PVDF membrane (Invitrogen). In order to probe *KRAS* and *β-actin* separately, the membrane was cut into two pieces according to the protein ladder and blocked with 5% non-fat milk. Then, the membranes were incubated with anti-KRAS antibody (Abcam 55391) and anti-β-actin antibody (Sigma AC-40) at 4°C overnight, respectively, followed by washing and incubating with secondary antibody (IRDye 800 CW, LI-COR) at room temperature for 2 hours. Finally, the protein bands were visualized using Odyssey Clx imaging system (LI-COR). Densitometry measurements were calculated using the gel analysis tool in ImageJ.

To evaluate the intracellular uptake of nanoparticles in cells with decreased *KRAS* expression, MDA-MB-231 cells were seeded in 24-well plates and transfected with 1.0 μM and 1.5 μM Accell siRNAs, respectively. 120 hours after transfection, cells were incubated with 500 μg/mL FITC-NP for 30 minutes and subsequently analyzed by flow cytometry using the same method as described above.

### Statistical Analysis

All experiments were performed in triplicate at minimum. The results were expressed as means ± standard deviation. Statistical significance was analyzed using Students’ t-test for mean differences among the samples.

## Results

### Physicochemical properties of nanoparticles

Cross-linked albumin nanoparticles were synthesized by desolvation method ^24,25^. The hydrodynamic size and charge of the nanoparticles were characterized by dynamic light scattering and zeta potential measurements, respectively. The sizes of the nanoparticles were controlled by changing the ratio of methanol and ethanol in the desolvation agent. As shown in Figure 1A, increasing the percentage of ethanol from 0 % to 100 % increased the mean diameter of the nanoparticles from 36.13 ± 0.27 nm to 252 ± 0.172 nm. The polydispersity indexes of the nanoparticles were all below 0.25, which suggest that the synthesis resulted in relatively monodisperse populations. The zeta potentials of the nanoparticles ranged from −37.60 ± 0.53 mV to −46.67 ± 0.32 mV. There is no obvious trend correlating particle size and charge with the change of desolvation agent composition (Figure 1B). The nanoparticles synthesized with desolvation agent of methanol: ethanol at the ratio of 3:7 (v/v) were chosen for subsequent experiments. The mean size of cross-linked nanoparticles with this ratio was 69.78 ± 0.43 nm with PDI of 0.13 ± 0.01. The zeta potential of the nanoparticles was −42.73 ± 1.20mV.

**Figure 1.**
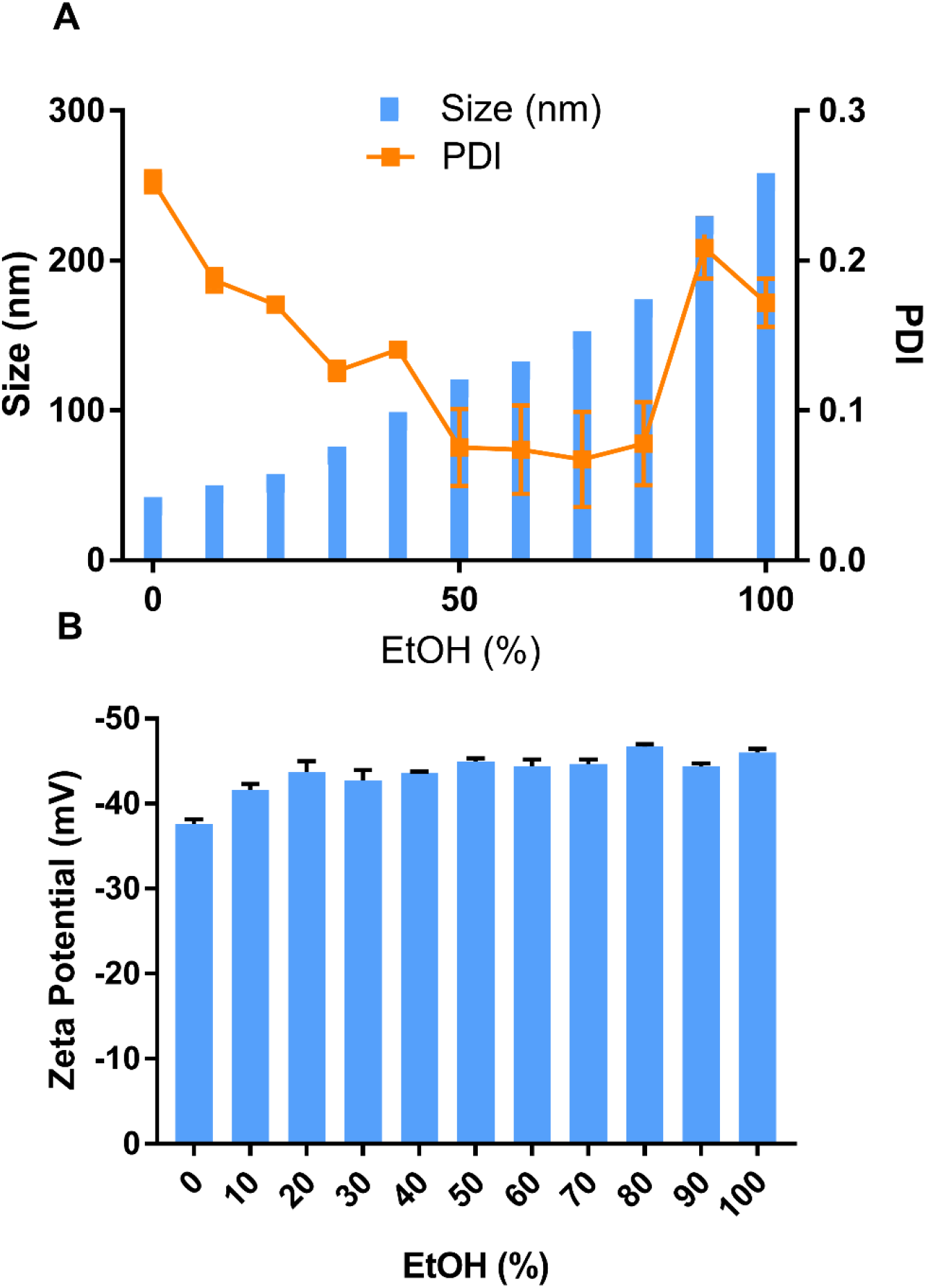
(A) Size and polydispersity index of nanoparticles synthesized at different EtOH (%). (B) Zeta potential of nanoparticles synthesized at different EtOH (%). (n = 3)

For subsequent studies to investigate cell uptake of particles, cross-linked albumin nanoparticles (FITC-NPs) were prepared using fluorescently-labeled monomeric albumin to ensure cross-linked nanoparticles had equivalent amount of FITC per albumin. The mean size and zeta potential of the FITC-NPs were 71.41 ± 0.64 nm (PDI 0.10 ± 0.004) and −42.5 ± 0.36 mV, respectively. These measurements indicate that the conjugation of FITC to albumin prior to nanoprecipitation of cross-linked NPs did not impact their physicochemical properties. Both non-labeled and FITC-labeled NPs were observed by transmission electron microscopy (Figure 2A and Figure 2B, respectively).

**Figure 2.**
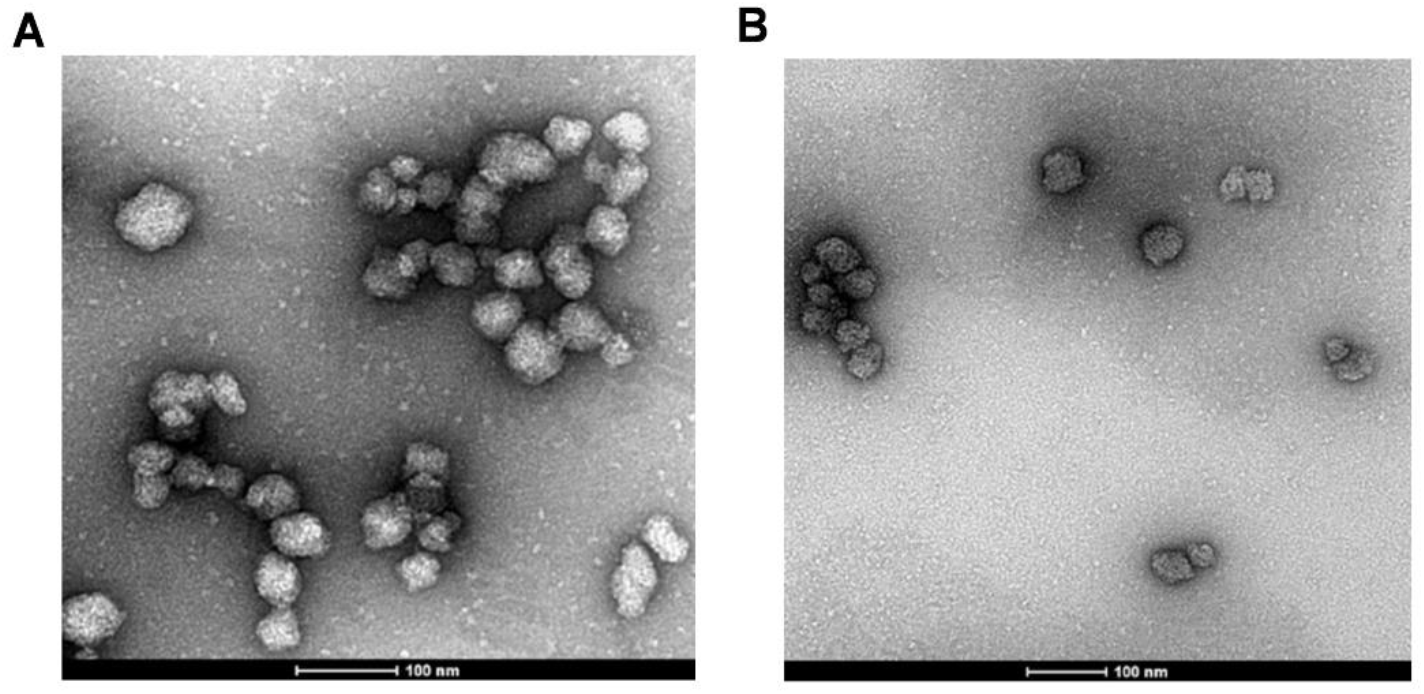
TEM images of (A) albumin nanoparticles and (B) FITC labelled nanoparticles.

The nanoparticles had a spherical morphology and were evenly distributed. To confirm their physiological stability *in vitro*, particles were incubated in complete media with 10% FBS at 37°C. The size and PDI of nanoparticles (Figure 3A and 3B, respectively) had negligible change up to 5 days, which confirms their serum stability.

**Figure 3.**
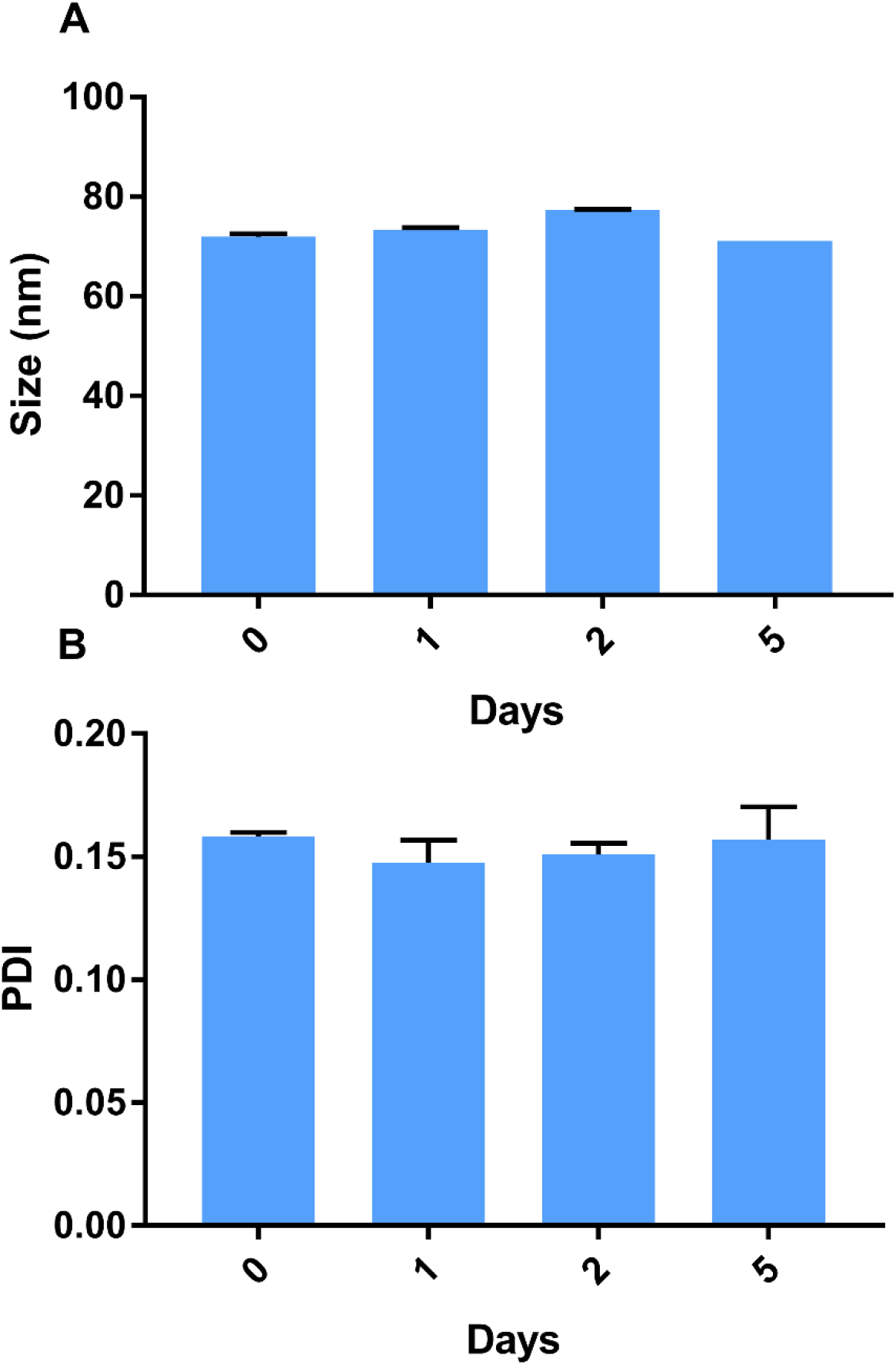
Stability of NPs incubated with 10% FBS DMEM, (A) size and (B) PDI measured at predetermined time-points. (n = 3)

### Nanoparticles demonstrate higher uptake than monomeric albumin

Intracellular uptake of FITC-BSA and FITC-NPs was evaluated in MDA-MB-231 cells with oncogenic *KRAS* mutation G13D (Figure 4A) and control MDA-MB-468 cells with wild-type *KRAS* alleles (Figure 4B). Equivalent amounts of FITC-BSA and FITC-NPs were incubated, and uptake of these fluorescent particles was quantified by flow cytometry. As shown in Figure 4, the mean fluorescence intensity (MFI) of FITC-BSA and FITC-NPs in both cell lines increased in a dose-dependent manner. However, at each dose, the MFI of cells treated by FITC-NPs was significantly higher than those of cells treated by FITC-BSA, which indicates that cross-linked NPs demonstrate greater uptake than monomeric albumin.

**Figure 4.**
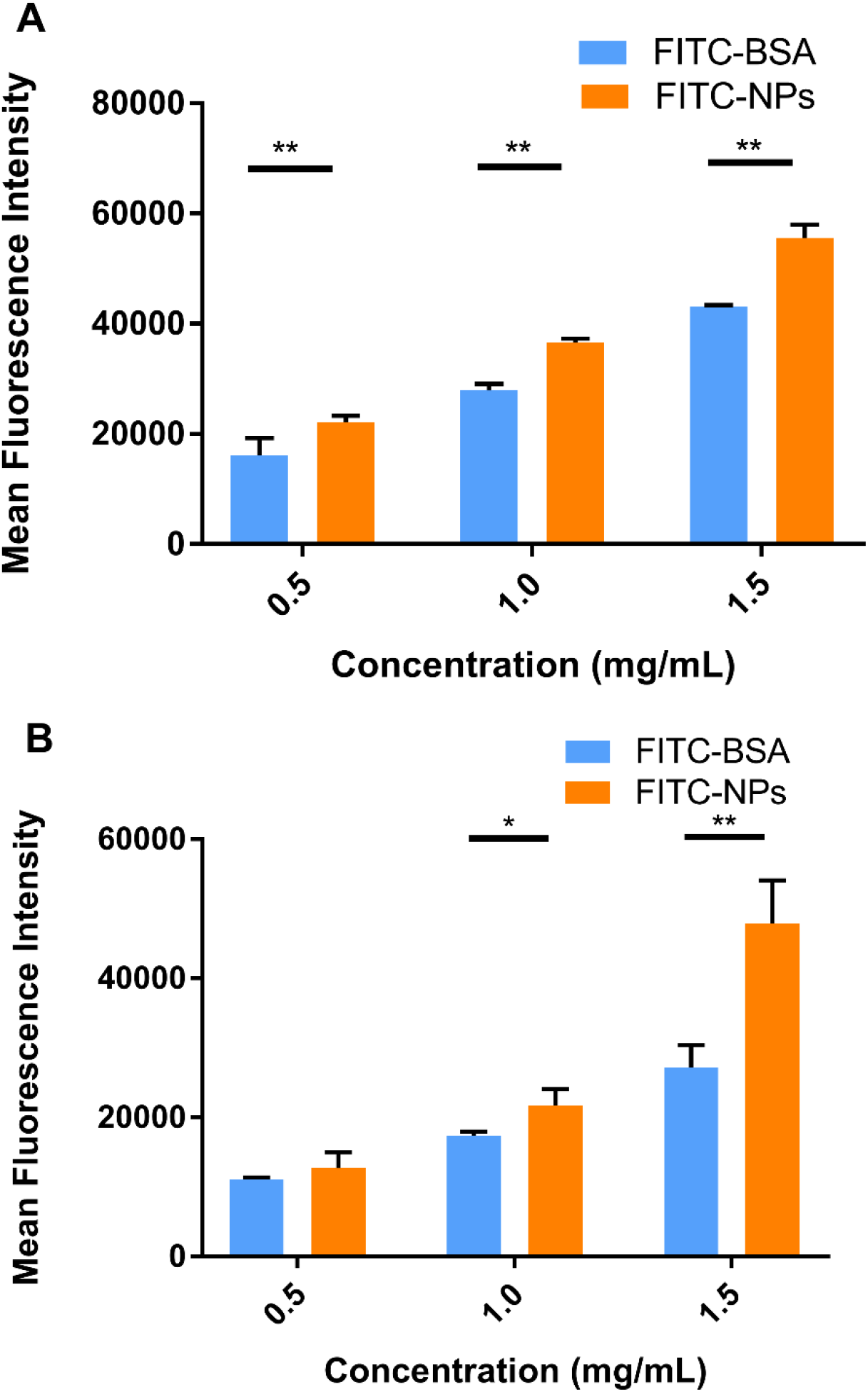
In vitro uptake of FITC-BSA and FITC-NPs in (A) MDA-MB-231 and (B) MDA-MB-468 cells. (n = 3, * *p* < 0.05, ** *p* < 0.01)

Cross-linked albumin may stabilize and improve upon existing *nab-paclitaxel;* upon systemic administration, *nab-paclitaxel* dissociates from particulate form into monomeric albumin and thereby limits the amount of drug delivery without incurring systemic toxicities ^18,20,26^. From the MFI values, oncogenic *KRAS* MDA-MB-231 cells exhibited greater uptake of both monomeric albumin and cross-linked albumin NPs than control MDA-MB-468 cells. Subsequent experiments were performed to support that activating mutations of *RAS* stimulate greater particle uptake than in cells with wild-type *KRAS* alleles.

### Decreased KRAS protein expression resulted in reduced intracellular uptake of nanoparticles

It was next tested if activating *KRAS* stimulates nanoparticle uptake in cancer cells. MDA-MB-231 cells were transfected with siRNA targeting *KRAS*, and knockdown resulted in decreased *KRAS* protein expression, as indicated by immunoblotting (Figure 5A). As shown in Figure 5A, siRNA knockdown of *KRAS* in cells at different concentrations decreased *KRAS* expression compared to non-targeting siRNA treated cells. Using densitometry to semi-quantify protein expression, *KRAS* protein expression decreased to 50.4% with 1.5 μM siRNA treatment. Subsequently, the intracellular uptake of FITC-NPs was determined in MDA-MB-231 cells with siRNA-mediated knockdown of *KRAS*. As shown in Figure 5B, there was no difference between cells without any treatment and cells treated with non-targeting siRNA. The addition of control siRNA did not affect the uptake of FITC-NPs. However, the uptake of FITC-NPs was significantly decreased in cells treated with *KRAS* targeting siRNAs, which indicates that direct inhibition of *KRAS* can negatively impact uptake of FITC-NPs. This finding supports previous studies that demonstrated hyperactivating *RAS* in cells stimulates their uptake of macromolecules ^10,11,13,14^.

**Figure 5.**
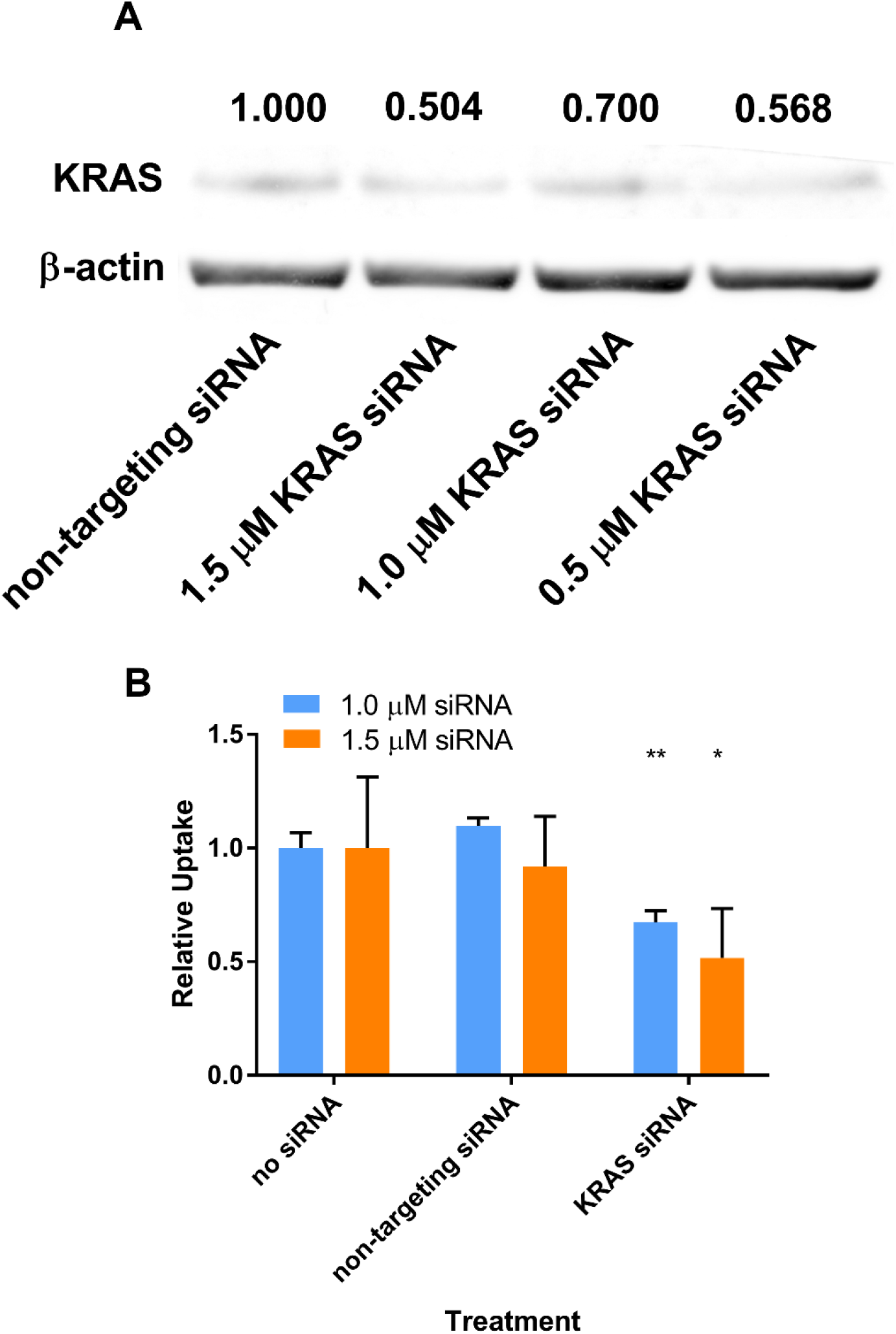
(A) Protein expression of *KRAS* and *β-actin* in MDA-MB-231 cells harvested 120 hours post-transfection by siRNA. Normalized values of the densitometric measurements are listed in the figure. (B) Relative uptake of FITC-NPs by MDA-MB-231 cells 120 hours post-transfection by siRNA. (n = 3, * *p* < 0.05, ** *p* < 0.01)

### Elevated macropinocytosis of nanoparticle uptake in oncogenic KRAS cells

It was then confirmed that oncogenic *KRAS* MDA-MB-231 exhibited increased macropinocytosis compared to cells with wild-type *RAS*. The macropinocytic activity of MDA-MB-231 cells and MDA-MB-468 cells was visualized by imaging the uptake of tetramethylrhodamine (TMR)-dextran, a fluorescent tracer for macropinocytosis. Here, TMR-dextran was internalized into cells via macropinosomes, shown as red puncta in Figure S1 (supplementary data). The amount of macropinosomes, which correlates with the extent of macropinocytosis, was quantified by calculating their macropinocytic index, as developed by Commisso et al. ^27^. As shown in Figure S1, the relative uptake of TMR-dextran in MDA-MB-231 cells was 2.5-fold higher than in MDA-MB-468 cells (p<0.01). Greater uptake of TMR-dextran in oncogenic *KRAS* cells was due to macropinocytosis, as confirmed with pharmacological inhibition of EIPA, a canonical inhibitor of macropinocytosis (Figure S1). After treatment with 25 μM EIPA, the relative macropinocytic index of MDA-MB-231 cells significantly decreased (p<0.01). However, there was no statistical difference in TMR-dextran uptake of MDA-MB-468 cells with or without EIPA (Figure S1). These results confirm that mutant *KRAS* cancer cells exhibit greater macropinocytosis than cells with wild-type *KRAS* alleles.

After confirming oncogenic *KRAS* MDA-MB-231 macropinocytoses reporter tracer TMR-dextran, it was next confirmed that the uptake of our cross-linked albumin nanoparticles is inhibited by a similar mechanism. As shown in Figure 6A, when MDA-MB-231 cells were treated with 25, 50, and 75 μM EIPA, the uptake of FITC-NPs were significantly inhibited by 16.79%, 21.50% and 16.03%, respectively. For MDA-MB-468 cells (Figure 6B), the inhibition ratios were 6.09%, 9.39%, and 18.40%, respectively. A larger percentage of FITC-NPs was inhibited by EIPA in MDA-MB-231 cells harboring oncogenic *KRAS* mutation compared to MDA-MB-468 cells with wild-type *KRAS*. The decrease in uptake due to the EIPA inhibition indicates that cross-linked albumin NPs can be endocytosed by macropinocytosis, and this decrease is more pronounced in mutant *KRAS* cells; this finding is comparable to other reports demonstrating increased albumin uptake by oncogenic *KRAS* cancer cells and tumors via macropinocytosis^7,11,12^.

**Figure 6.**
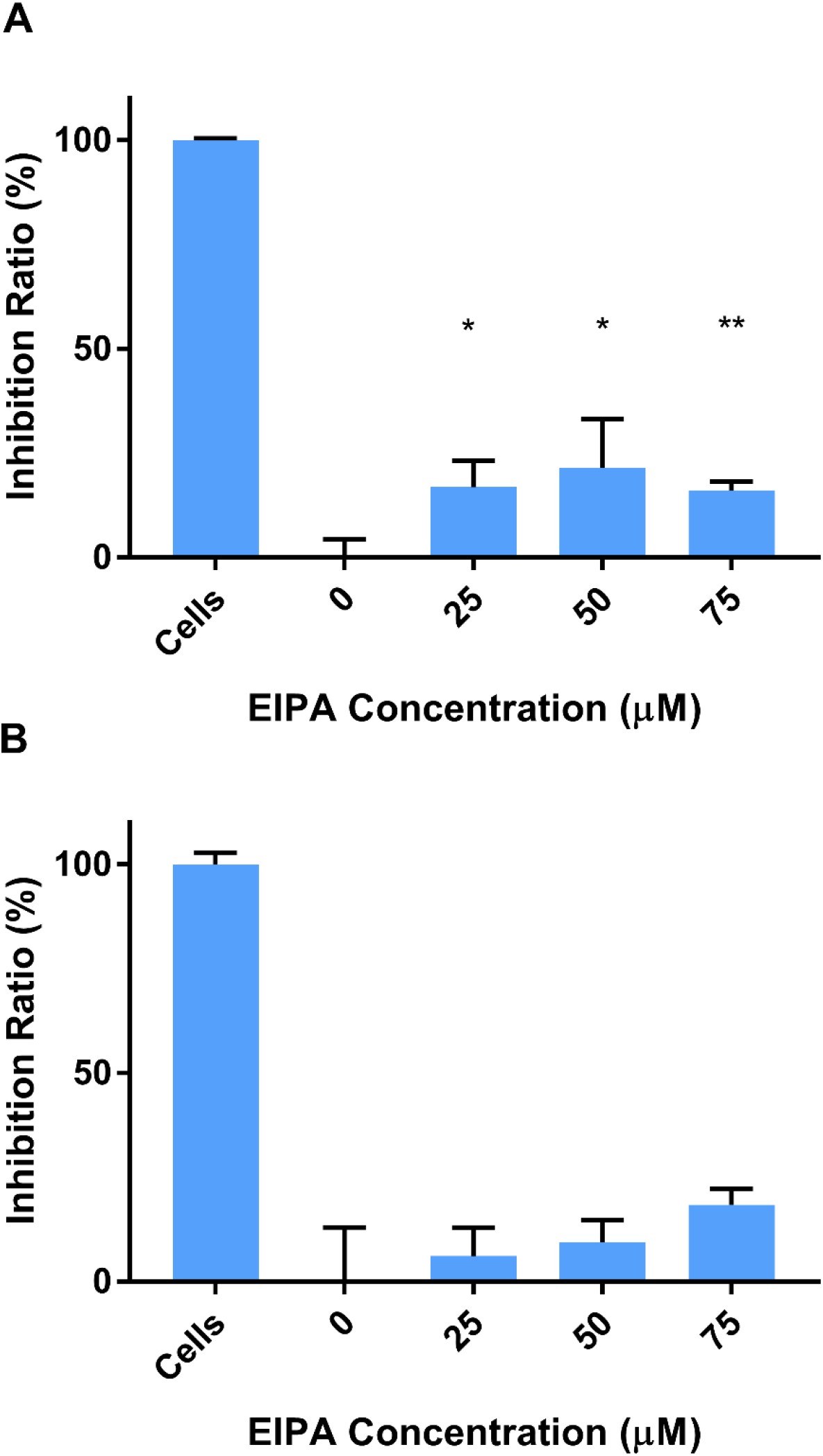
EIPA inhibited uptake of FITC-NPs in (A) MDA-MB-231 cells and (B) MDA-MB-468 cells. (n = 3, * *p* < 0.05, ** *p* < 0.01)

### Colocalization of nanoparticles with macropinosomes in oncogenic KRAS cells

To further confirm that albumin NPs were present in macropinosomes, which are indicative of macropinocytic uptake, Cy7 labelled nanoparticles were synthesized for the colocalization analysis. The hydrodynamic size and zeta potential of Cy7 nanoparticles were 62.22 ± 0.55 nm and −45.70 ± 0.42 mV. Colocalization of Cy7 nanoparticles with TMR-dextran was shown in Figure 7. As shown in Figure 7A, Cy7 nanoparticles (green) and TMR-dextran (red) were both taken up by mutant *KRAS* MDA-MB-231 cells. After treatment with EIPA (Figure 7B), the amount of red and green puncta both decreased, indicating the uptake of TMR-dextran and Cy7 nanoparticles was decreased. In Figure 7C and 7D, the uptake level of Cy7 nanoparticles and TMR-dextran were similar in MDA-MB-468 cells, regardless of the EIPA inhibition. To determine the localization of albumin NPs in macropinosomes, we quantitatively correlated the co-localization of NPs with TMR-dextran marker via image analysis and calculation of the Pearson Correlation Coefficient (PCC). The PCC values between the two channels were summarized in Figure 7E. The PCC value for MDA-MB-231 cells was 0.82, which indicates good correlation of pixel intensity distribution between red and green channels. After EIPA inhibition, the PCC value for MDA-MB-231 significantly decreased to 0.49 (p<0.05), which means there was a significant decrease of nanoparticle uptake by macropinocytosis. The relative low PCC values in MDA-MB-468 cells indicated that the Cy7 nanoparticles did not colocalize well with TMR-dextran as those in MDA-MB-231 cells. In other words, the fraction of nanoparticles taken up by macropinocytosis was lower than that in oncogenic *KRAS* MDA-MB-231 cells.

**Figure 7.**
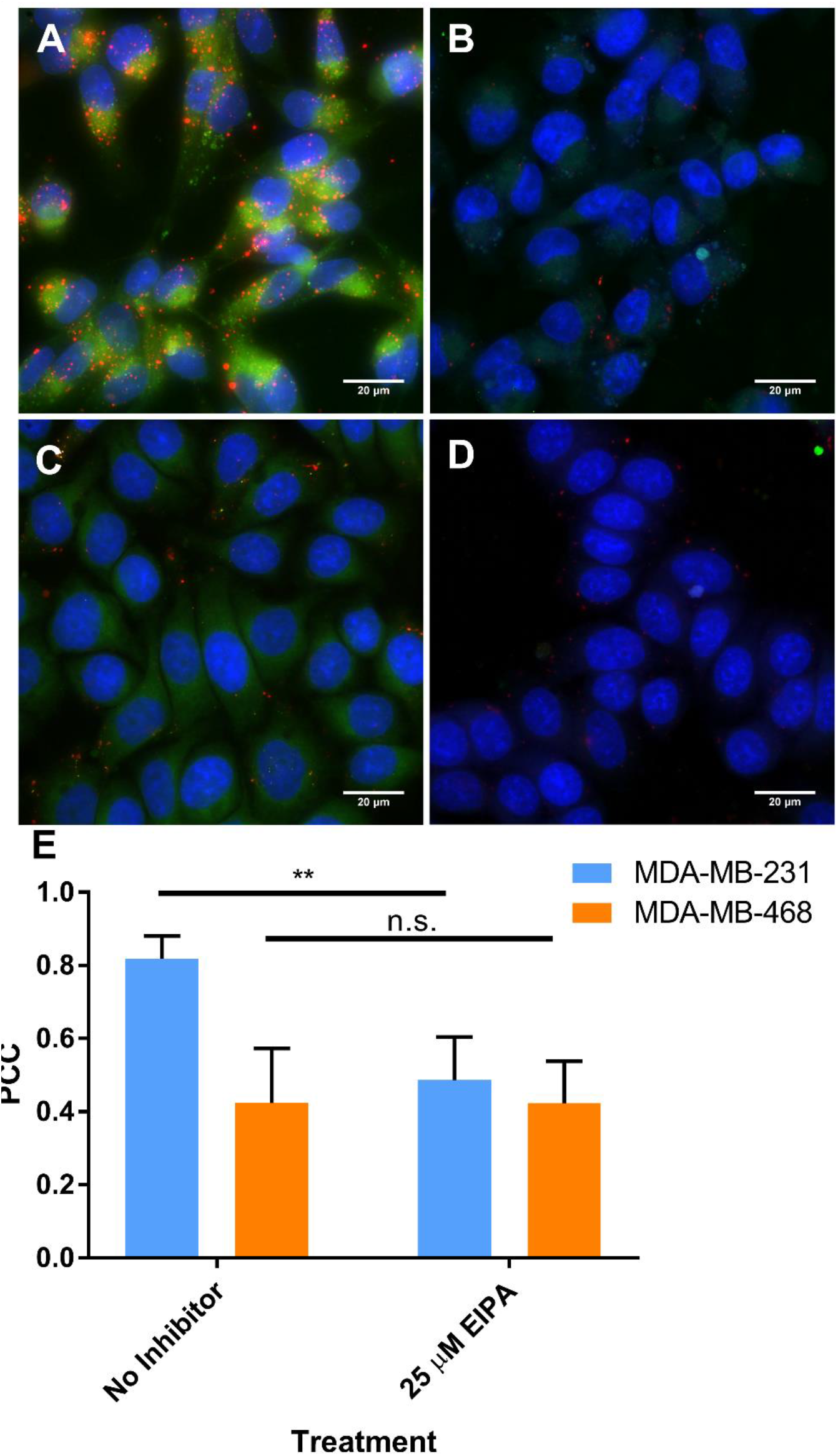
Extended focus images of intracellular uptake of TMR-dextran (red) and Cy7-NPs (green): (A) MDA-MB-231 cells; (B) MDA-MB-231 cells treated with 25 μM EIPA; (C) MDA-MB-468 cells; (D) MDA-MB-468 cells treated with 25 μM EIPA. (E) Pearson correlation coefficient of Cy7-NPs vs. TMR-dextran in MDA-MB-231 and MDA-MB-468 cells. (n = 10, ** *p* < 0.01, *n.s*. is no significant difference)

## Discussions

Oncogenic *RAS*, the most abundant overall mutation in cancers, programs cell signaling pathways and tumor progression; however, drugging this difficult-to-target oncoprotein has been the promise and a long-standing goal in cancer therapy. Hyperactive *RAS* triggers intake of extracellular nutrients needed for biogenesis, cancer cell survival, and proliferation to worsen tumor progression. Studies demonstrated that mice with pancreatic tumors possessing *KRAS* mutations demonstrated greater uptake of radiolabeled mouse serum albumin than healthy pancreas and wild-type KRAS pancreatic cancers ^11,12^. This increased uptake of albumin has also been previously observed with RAS-transformed fibroblasts ^10,14^, glioblastoma ^28^. Further, it was found that scavenged and catabolized extracellular albumin was the only source of amino acids present in oncogenic *KRAS* tumors and was needed for cancer cell proliferation and DNA synthesis ^12^. Davidson et al. confirmed that albumin was macropinocytosed by oncogenic *KRAS* tumors and then degraded in lysosomes *in vivo* ^12^. In addition to albumin and lipid uptake by oncogenic *KRAS* cells via macropinocytosis ^11–14,17^, it has been recently demonstrated that larger-sized solutes, including antibodies ^21^ and lipoprotein nanoparticles ^20^, also exhibit increased uptake in active RAS-stimulated cells via micropinocytosis. Constitutively active *RAS* stimulates intracellular uptake of lipids and proteins through the endocytic route of macropinocytosis, which facilitates transport of large solutes in vesicles up to 5 μm in diameter. Leveraging the metabolic needs of cancers for solute uptake via macropinocytosis, we wanted to develop nanoparticles that specifically target this mechanism present in RAS-transformed or oncogenic cancer cells towards drug delivery. Building on prior studies demonstrating albumin uptake and catabolism in mutant *KRAS* cancers, we synthesized stable albumin nanoparticles and tested their ability to enter oncogenic *KRAS* cancer cells for intracellular delivery. We confirmed that albumin NPs exhibit significantly enhanced uptake in oncogenic *KRAS* cancer cells compared with control cells with wild-type *KRAS*. Through extensive pharmacological inhibition, genetic knockdown, and microscopy studies, we demonstrated hyperactivated *KRAS* is responsible for stimulating macropinocytosis to engulf albumin NPs. By targeting oncogenic RAS-driven macropinocytosis for delivery, there is no need for conjugated ligands on the drug delivery system to facilitate cell binding and internalization. This simplicity of the nanoprecipitation synthesis and lack of conjugation chemistries avoids the challenges of chemistry and scalability and highlights the potential of this carrier for translation ^29,30^. While future studies to confirm cell specificity are needed, these albumin nanoparticles exhibit improved uptake in mutant *RAS* cancer cells compared to controls. Interestingly, albumin NPs demonstrated significantly improved uptake compared to equivalent amount of albumin monomers in cancer cell lines and in particular oncogenic *KRAS* MDA-MB-231 cells. The current gold standard in albumin drug carriers, *nab-paclitaxel*, is a 130 nm paclitaxel-loaded albumin nanoparticle in formulation ^31^; however, upon systemic administration, the particle rapidly dissociates into albumin monomers equivalent to endogenous albumin, and the drug prematurely releases, resulting in promiscuous accumulation in non-tumor tissues and organs and off-target toxicities ^18,20,26^. Coupled with the information that the cross-linked albumin NPs are stable in serum for several days, these findings suggest that cross-linked albumin NPs could deliver a higher amount of albumin and potentially, drug, than non-covalent, monomeric albumin-associated *nab-paclitaxel*. Additional studies will be needed to confirm drug encapsulation and stability of the cross-linked particles and compare its efficacy to *nab-paclitaxel* in cell culture and tumors. Finally, these nanoparticles were formulated to actively target the macropinocytic pathway for intracellular uptake in oncogenic *RAS* cancers; to fulfill the promise of these nanoparticles for RAS-targeted therapy, it will be necessary to extend delivery to additional cancers with more canonical *RAS* mutations (e.g. *KRAS* G12V, G12C; *NRAS* G12D) than the cells used in this current study and to use RAS-targeted therapeutics such as covalent inhibitors ^32,33^, stapled peptides ^34^, and other downstream Raf/MEK/ERK inhibitors ^1^.

## Conclusions

In this work, the synthesized cross-linked albumin nanoparticles exhibit significant uptake in oncogenic KRAS cancer cells compared with control wild-type KRAS cells. Oncogenic KRAS mutation is responsible for driving macropinocytosis to engulf albumin nanoparticles. This initial step to use stable albumin-based particles to exploit the metabolic vulnerability of oncogenic *RAS* for intracellular uptake opens new avenues for macropinocytic-driven drug delivery targeting the formidable barrier of oncogenic RAS-driven cancers.

