## Supplemental FIgure S1 for "Intracellular nanoparticle delivery by oncogenic KRAS-mediated macropinocytosis"

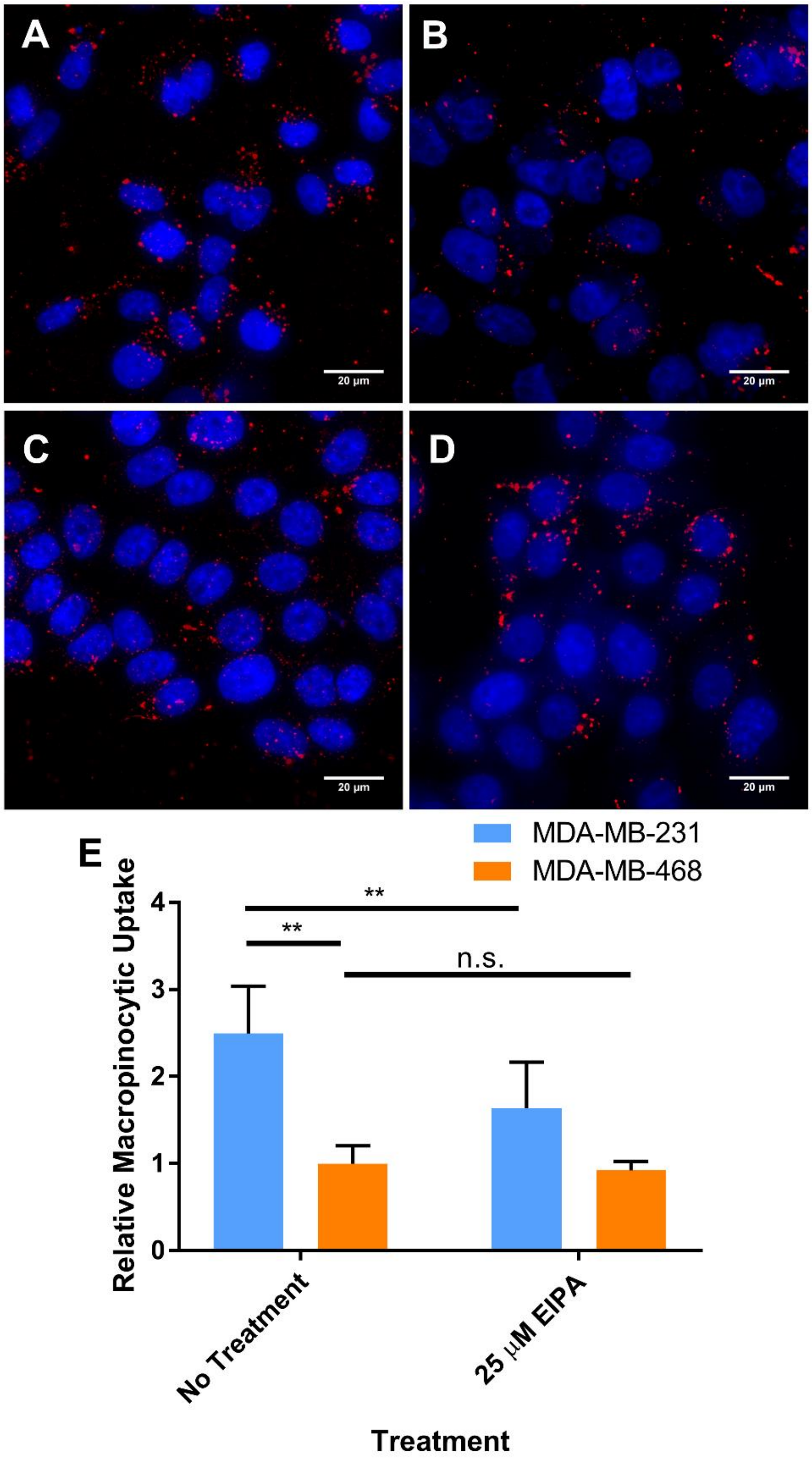


Figure S1. Macropinocytosis of TMR-dextran (red) in cells (nucleis in blue) observed by fluorescence microscopy. (A) MDA-MB-231 cells without EIPA treatment; (B) MDA-MB-231 cells treated with 25 µM EIPA; (C) MDA-MB-468 cells without EIPA treatment; (D)

MDA-MB-468 cells treated with 25 µM EIPA. (D) Relative macropinocytotic index of cells. ($n = 10$, ** $p < 0.05$, n.s. is no significant difference)
